# Minimizer-space de Bruijn graphs

**DOI:** 10.1101/2021.06.09.447586

**Authors:** Barış Ekim, Bonnie Berger, Rayan Chikhi

## Abstract

DNA sequencing data continues to progress towards longer reads with increasingly lower sequencing error rates. We focus on the problem of assembling such reads into genomes, which poses challenges in terms of accuracy and computational resources when using cutting-edge assembly approaches, e.g. those based on overlapping reads using minimizer sketches. Here, we introduce the concept of *minimizer-space* sequencing data analysis, where the minimizers rather than DNA nucleotides are the atomic tokens of the alphabet. By projecting DNA sequences into ordered lists of minimizers, our key idea is to enumerate what we call *k-min-mers*, that are k-mers over a larger alphabet consisting of minimizer tokens. Our approach, mdBG or minimizer-dBG, achieves orders-of-magnitude improvement in both speed and memory usage over existing methods without much loss of accuracy. We demonstrate three uses cases of mdBG: human genome assembly, metagenome assembly, and the representation of large pangenomes. For assembly, we implemented mdBG in software we call `rust-mdbg`, resulting in ultra-fast, low memory and highly-contiguous assembly of PacBio HiFi reads. A human genome is assembled in under 10 minutes using 8 cores and 10 GB RAM, and 60 Gbp of metagenome reads are assembled in 4 minutes using 1 GB RAM. For pangenome graphs, we newly allow a graphical representation of a collection of 661,405 bacterial genomes as an mdBG and successfully search it (in minimizer-space) for anti-microbial resistance (AMR) genes. We expect our advances to be essential to sequence analysis, given the rise of long-read sequencing in genomics, metagenomics and pangenomics.

## Introduction

DNA sequencing data continues to improve from long reads of poor quality [2], used to assemble the first human genomes, to Illumina short reads with low error rates (≤ 1%) and now to longer reads with low error rates. For instance, recent Pacific Biosciences (PacBio) instruments can sequence 10 to 25 Kbp-long (HiFi) reads at ≤ 1% error rate [52]. The R10.3 pore of Oxford Nanopore produces reads of hundreds of Kbps in length at a ~ 5% error rate. A tantalizing possibility is that DNA sequencing will eventually converge to long, nearly-perfect reads. These new technologies require novel algorithms that are both efficient and accurate for important sequence analysis tasks such as genome assembly [34].

Efficient algorithms for sequence analysis have played a central role in the era of high-throughput DNA sequencing. Many analyses such as read mapping (e.g. [54,50]), genome assembly (e.g. [44]), and taxonomic profiling (e.g., [37,40]) have benefited from milestone advances that effectively compress, or sketch, the data [35]; e.g. fast full-text search with the Burrows-Wheeler transform (BWT) [8], space-efficient graph representations with succinct de Bruijn graphs [11], and lightweight databases with MinHash sketches [42]. Large-scale data re-analysis initiatives [18, 27] further incentivize the development of efficient algorithms, as they aim to re-analyze petabases of existing public data.

However, there has traditionally been a tradeoff between algorithmic efficiency and loss of information, at least during the initial sequence processing steps. Consider short-read genome assembly: The non-trivial insight of chopping up reads into *k*-mers, thereby bypassing the ordering of *k*-mers within each read, has unlocked fast and memory-efficient approaches using de Bruijn graphs; yet the short *k*-mers—chosen for efficiency—lead to fragmented assemblies [3]. In modern sequence similarity estimation and read mapping approaches [54] information loss is even more drastic as large genomic windows are sketched down to comparatively tiny sets of *minimizers*—which index a sequence (window) by its lexicographically smallest *k*-mer [42], and enable efficient but sometimes inaccurate comparisons between gigabase-scale sets of sequences [23].

Here, we provide a highly efficient genome assembly tool for state-of-the-art and low-error long read data. We introduce minimizer de Bruijn graphs, mdBG, which instead of building an assembly over sequence bases—the standard approach that for clarity we refer to as *base-space*—newly performs assembly in *minimizer-space* (Figure 1), that is later converted back to base-space assemblies. Specifically, each read is initially converted to an ordered sequence of its minimizers [47, 32]. The order of the minimizers is important as our aim is to reconstruct the entire genome as an ordered list. This is in contrast with the classical MinHash technique, which converts sequences into unordered sets of minimizers to detect pairwise similarities between them [7]. To aid in assembly of higher-error rate data, we also introduce a novel variant of the partial order alignment (POA) algorithm, that operates in minimizer-space instead of base-space, and effectively corrects only the bases corresponding to minimizers in the reads. Sequencing errors that occur outside minimizers do not affect our representation; those within minimizers cause substitutions or indels in minimizer-space (Figure 2), which can be identified and subsequently corrected in minimizer-space using POA (Figure 3).

**Fig. 1:**
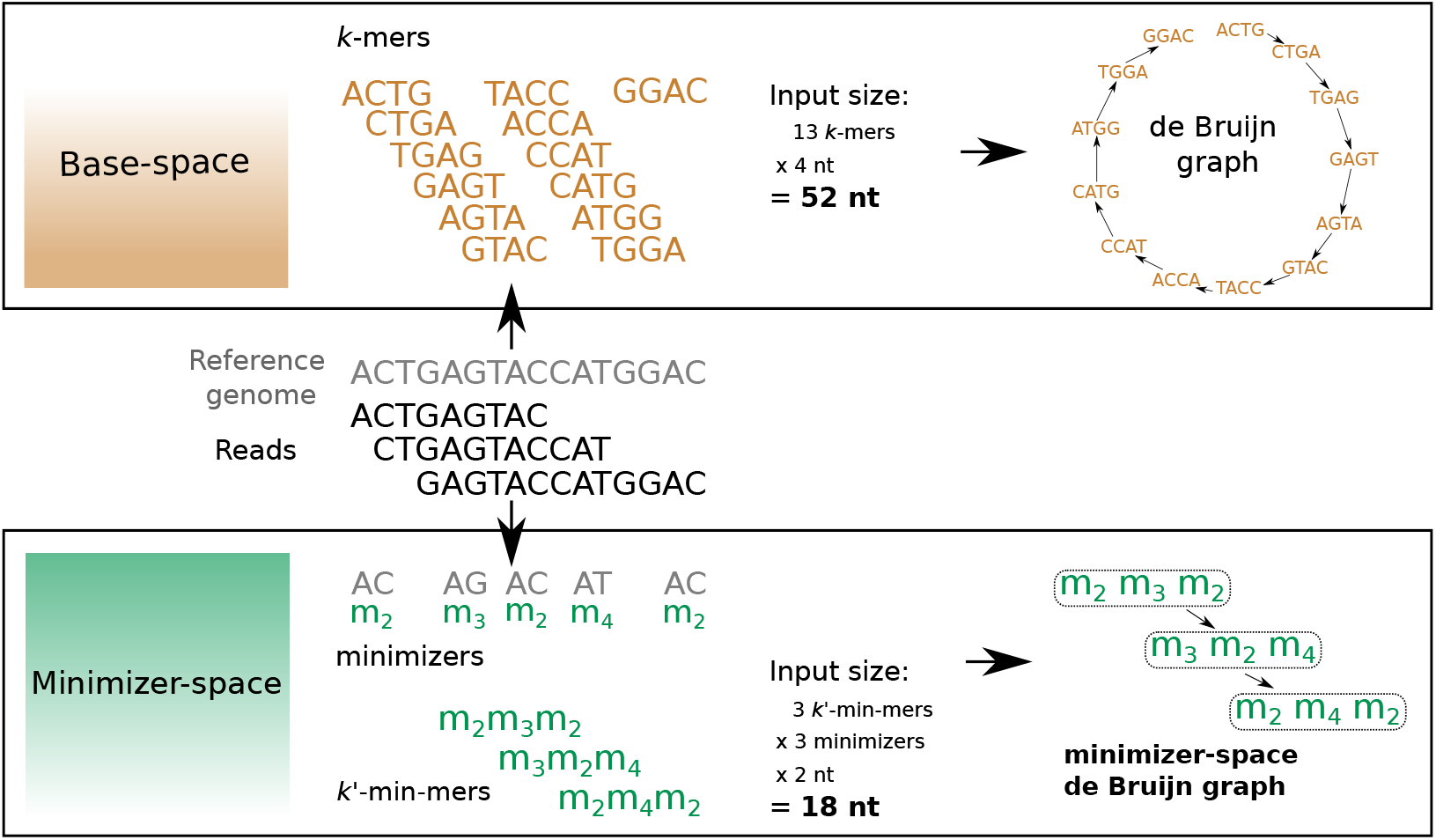
An efficient assembly method for state-of-the-art genome sequencing (e.g. PacBio HiFi data). Illustration of our minimizer-space de Bruijn graph (mdBG, bottom) compared to the original de Bruijn graph (top). Center horizontal section shows a toy reference genome, along with a collection of sequencing reads. Top box shows *k*-mers (*k* = 4) collected from the reads, which are the nodes of the classical de Bruijn graph. The input size of 52 nucleotides (nt) is depicted in boldface. Bottom box shows the position of minimizers in the reads for *ℓ* = 2, and any *ℓ*-mer starting with nucleotide ‘A’ is chosen as a minimizer. *k*’-min-mers (*k*’ = 3) are tuples of *k*’ minimizers as ordered in reads, which constitute the nodes of the minimizer-space de Bruijn graph. The reduction in input size to 18 nucleotides (nt) is depicted in boldface.

**Fig. 2:**
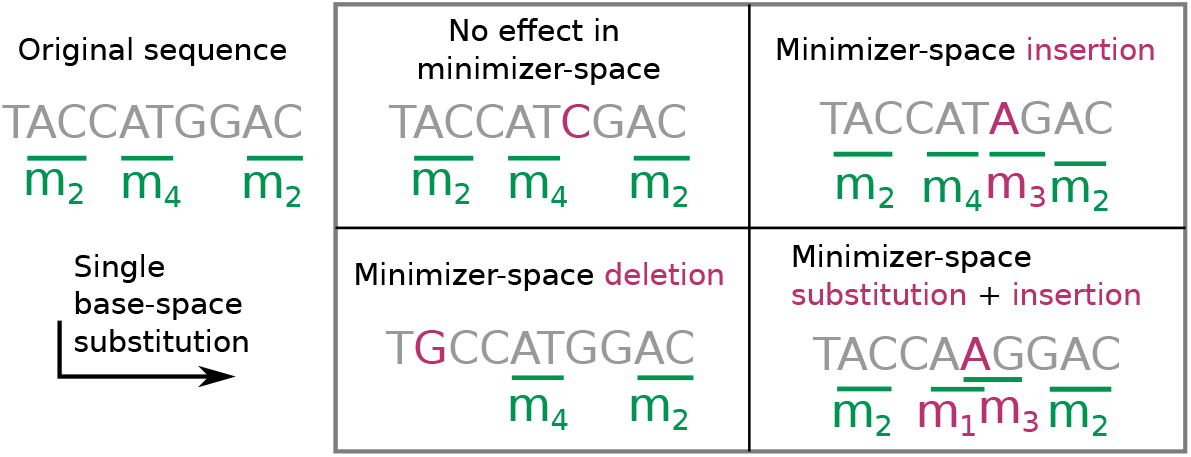
Propagation of sequencing errors in base-space to minimizer-space. Continuing the example in Figure 1, we consider a sequence along with its minimizers (left of the box). Each panel inside the box depicts the effect of a different mutation on the sequence. Top left panel: G→C (in purple) leads to no change in the minimizer-space representation as the mutation did not change or create any minimizer. Bottom left: A→G led to the disappearance of *m*_2_. Top right: C→A made the *m*_3_ minimizer appear. Bottom right: T→A affected two minimizers: *m*_4_ was substituted for *m*_1_, and *m*_3_ was inserted.

**Fig. 3:**
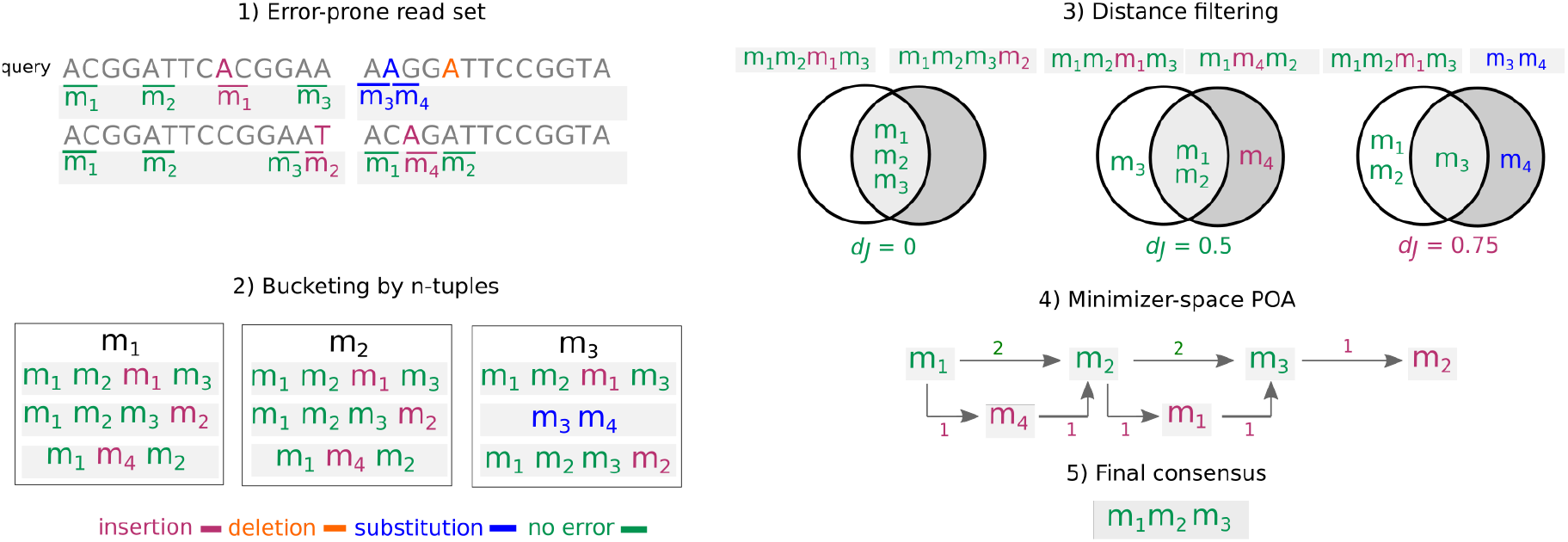
Overview of the minimizer-space partial order alignment (POA) procedure. **1)**with a toy dataset of 4 reads. Error-prone reads and their ordered lists of minimizers (*ℓ* = 2) are shown, with sequencing errors and the minimizers that are created as a result of errors denoted in colors (insertion as red, deletion as orange, substitution in blue, no errors in green). **2)** Before minimizer-space error-correction, the ordered lists of minimizers are bucketed using their *n*-tuples (*n* = 1). **3)** For a query ordered list (the first read in the read set in the figure), all ordered lists that share an *n*-tuple with the query are obtained, and the final list of query neighbors are obtained by applying a heuristically determined distance filter *d_j_* (Jaccard distance threshold of *φ* = 0.5). **4)** A POA graph in minimizer-space is constructed by initializing the graph with the query, and aligning each ordered list that passed the filter to the graph iteratively (weights of poorly-supported edges are shown in red). **5)** By taking a consensus path of the graph, the error in the query is corrected.

Our key conceptual advance is that minimizers can themselves make up atomic tokens of an extended alphabet, which enables efficient long-read assembly that, along with error correction, leads to preserved accuracy. We can instead perform assembly using a minimizer-space de Bruijn graph (mdBG), drastically reducing the amount of data input to the assembler, preserving accuracy, lowering running time, and decreasing memory usage by 1 to 2 orders of magnitude compared to current assemblers. Setting adequate parameters for the order of the de Bruijn graph and the density of our minimizer scheme allows us to overcome stochastic variations in sequencing depth and read length, in a similar fashion to traditional base-space assembly. To handle higher sequencing error rates, we newly correct for base errors by performing partial order alignment instead in minimizer-space.

With error-prone data, we study two regimes: Real PacBio HiFi read data (< 1% error rate) for *D. melanogaster* and Human, which turn out to require little adjustment for errors due to the very low rate, and synthetic 1 to 10% error rate data, which corresponds to the range of error rates of Oxford Nanopore’s recent technology. We also demonstrate that despite data reduction, running our `rust-mdbg` software on synthetic error-free and 4% error-rate data results in near-perfect reconstruction of a genome, the latter entirely due to our application of POA in minimizer-space.

To further demonstrate `rust-mdbg`’s capabilities, we use it to assemble two PacBio HiFi metagenomes, achieving runtimes of minutes as opposed to days and memory usage two orders of magnitude lower than the current state-of-the-art `hifiasm-meta`, with comparable assembly completeness yet lower contiguity. As a versatile use case of minimizer-space analysis, we construct to the best of our knowledge the largest pangenome graph to date of 661K bacterial genomes, and perform minimizer-space queries of antimicrobial resistance (AMR) genes within this graph, identifying nearly all those with high sequence similarity to original bacterial genomes. Rapidly detecting AMR genes in a large collection of samples would facilitate real-time AMR surveillance [20], and mdBG provides a space-efficient alternative to indexed *k*-mer search.

Remarkably, our approach is equivalent to examining a tunable fraction (e.g. only 1%) of the input bases in the data and should generalize to emerging sequencing technologies.

### Comparison with Related Work

This work is at the confluence of three core ideas that were just recently proposed in three different genome assemblers: `Shasta` [49], `wtdbg2` [48] and `Peregrine` [15]. 1) `Shasta` transforms ordered lists of reads into minimizers (`Shasta` used the term *markers*) to produce an efficiently reduced representation of sequences that facilitates quick detection of overlaps between reads. A similar idea was previously used for read mapping and assembly in `minimap/miniasm` [30, 31], and edit distance calculation with Order Min Hash (OMH) [38]. 2) The `wtdbg2` idea extends the usual *Σ* = {A,C,T,G} alphabet, which forms the basis of traditional genome de Bruijn graphs, to 256 bp windows: A ‘fuzzy’ de Bruijn graph is constructed by ‘zooming out’ of read sequences and considering batches of 256 base pairs (bps) at a time. 3) The `Peregrine` idea can be broken down into two parts: i) pairs of consecutive minimizers can be indexed, and they are naturally less often repeated across a genome than isolated minimizers; and ii) a hierarchy of minimizers can be constructed so that fewer minimizers are selected than in classical methods, thus increasing the distance between minimizers.

In distantly-related independent work, a very recent pre-print [46] (MBG) demonstrates a similar idea as `Peregrine`, performing assembly by finding pairs of consecutive minimizers on reads. Although MBG does combine the concepts of minimizers and de Bruijn graphs, it is fundamentally different from the work presented here: Nodes in the MBG are classical *k*-mers over the DNA alphabet, whereas nodes in our representation are *k*-mers over an alphabet of minimizers. Two other related concepts to MBG are sparse de Bruijn graphs [53] and A-Bruijn graphs [26, 33], in which the nodes are a subset of the original de Bruijn graph nodes, and the edge condition is relaxed so that overlaps may be shorter than (*k* — 1) when pairs of nodes are seen consecutively in a read.

Conceptually, our advance is in tightly combining both de Bruijn graphs and minimizers, a novel and non-trivial mix of previously-known ingredients. The concept of a de Bruijn graph was not considered in either the `Shasta` or the `Peregrine` assemblers; whereas in the `wtdbg2` assembler, de Bruijn graphs were considered, but not minimizers. Moreover, reducing the three aforementioned genome assemblers into a single idea for each of them, in terms of how they achieve algorithmic efficiency, is a contribution in itself and simplifies our presentation greatly. What we offer is essentially an ultra-fast variation of de Bruijn graphs, for long reads.

## Methods

### Resource and Materials Availability

This study did not generate new materials. Further information and requests for resources should be directed to and will be fulfilled by the Lead Contact, Rayan Chikhi. The code generated during this study is available at https://github.com/ekimb/rust-mdbg/.

### Method Details

#### Preliminaries

We encourage the reader to not skip the following definitions as they contain uncommon concepts. The variable *σ* is used as a placeholder for an unspecified *alphabet* (a non-empty set of *characters*). We define *Σ*_DNA_ = {A, C, T, G} as the alphabet containing the four DNA bases. Given an integer *ℓ* > 0, *Σ^ℓ^* is the alphabet consisting of all possible strings of length *ℓ*. To avoid confusion, we stress that *Σ^ℓ^* is an unusual alphabet: Any ‘character’ of *Σ^ℓ^* is itself a string of length *ℓ* over the DNA alphabet.

Given an alphabet *σ*, a *string* is a finite ordered list of characters from *σ*. Note that our strings will sometimes be on alphabets where each character cannot be represented by a single alphanumeric symbol. Given a string *x* over some alphabet *σ* and some integer *n* > 0, the *prefix* (respectively the *suffix*) of *x* of length *n* is the string formed by the first (respectively the last) *n* characters of *x*.

We now introduce the concept of a *minimizer*. In this paragraph we consider strings over the alphabet *Σ*_DNA_. We consider two type of minimizers: *universe* and *window*. Consider a function *f* that takes as input a string of length *ℓ* and outputs a numeric value within range [0, *H*], where *H* > 0. Usually, *f* is a 4-bit encoding of DNA or a random hash function (it does not matter whether the values of *f* are integers or whether *H* is an integer). Given an integer *ℓ* > 1 and a coefficient 0 < *δ* < 1, a *universe* (*ℓ, δ*)-*minimizer* is any string *m* of length *ℓ* such that *f*(*m*) < *δ · H*. We define *M_ℓ, δ_* to be the set of all universe (*ℓ, δ*)-minimizers, and we refer to *δ* as the *density* of *M_ℓ,δ_*.

This definition of a minimizer is in contrast with the classical one [47] which we recall here, although we will not use it. Consider a string *x* of any length, and a substring (*window*) *y* of length *w* of *x*. A window *ℓ-minimizer* of *x* given window *y* is a substring *m* of length *ℓ* of *y* that has the smallest value *f* (*m*) among all other such substrings in that window. Observe that universe minimizers are defined independently of a reference string, unlike window minimizers. They have been recently independently termed mincode syncmers [17]. We also performed experiments with an alternative concept to minimizers, Locally Consistent Parsing (LCP) [16], which replaces universal minimizers with *core substrings*: Substrings that can be pre-computed for any given alphabet such that any sequence of length *n* includes ~ *n*/*ℓ* substrings of length *ℓ* on average (Supplementary Note F).

We recall the definition of de Bruijn graphs. Given an alphabet *σ* and an integer *k* ≥ 2, a de Bruijn graph of order *k* is a directed graph where nodes are strings of length *k* over *σ* (*k*-mers), and two nodes *x,y* are linked by an edge if the suffix of *x* of length *k* — 1 is equal to the prefix of *y* of length *k* — 1. This definition corresponds to the node-centric de Bruijn graph [12] generalized to any alphabet.

#### Minimizer-space de Bruijn graphs

We say that an algorithm or a data structure operates in *minimizer-space* when its operations are done on strings over the *Σ^ℓ^* alphabet, with characters from *M_ℓ,δ_*. Conversely, it operates in *base-space* when the strings are over the usual DNA alphabet *Σ*_DNA_.

We introduce the concept of (*k, ℓ, δ*)-*min-mer*, or just *k-min-mer* when clear from the context, defined as an ordered list of *k* minimizers from *M_ℓ,δ_*. We use this term to avoid confusion with *k*-mers over the DNA alphabet. Indeed, a *k*-min-mer can be seen as a *k*-mer over the alphabet *Σ^ℓ^*, i.e. a *k*-mer in minimizer-space. For an integer *k* > 2 and an integer *ℓ* > 1, we define a *minimizer-space de Bruijn graph* (mdBG) of order *k* as de Bruijn graph of order *k* over the *Σ^ℓ^* alphabet. As per the definition in the previous section, nodes are *k*-min-mers, and edges correspond of identical suffix-prefix overlaps of length *k* — 1 between *k*-min-mers. Figure 1 shows an example.

We present our procedure for constructing mdBGs as follows. First, a set *M* of minimizers are pre-selected using the universe minimizer scheme from the previous section. Then, reads are scanned sequentially, and positions of elements in *M* are identified. A multiset *V* of *k*-min-mers is created by inserting all tuples of *k* successive elements in *M_ℓ,δ_* found in the reads into a hash table. Each of those tuples is a *k*-min-mer, i.e. a node of the mdBG. Edges of the mdBG are discovered through an index of all (*k* — 1)-min-mers present in the *k*-min-mers.

mdBGs can be simplified and compacted similarly to base-space de Bruijn graphs, using similar rules for removing likely artefactual nodes (tips and bubbles), and performing path compaction. They are also bidirected, though we present them as directed here for simplicity. See ‘Implementation details’ for more details on reverse complements and simplification.

By itself the mdBG is insufficient to fully reconstruct a genome in base-space, as in the best case it can only provide a sketch consisting of the ordered list of minimizers present in each chromosome. To reconstruct a genome in base-space, we associate to each *k*-min-mer the substring of a read corresponding to that *k*-min-mer. The substring likely contains base-space sequencing errors, which we address at the end of this paragraph. It is also necessary to keep track of the positions of the second and second-to-last minimizers in each *k*-min-mer. After performing compaction, the base sequence of a compacted mdBG can be reconstructed by concatenating the sequences associated to *k*-min-mers, making sure to discard overlaps. Note that in the presence of sequencing errors, or when the same *k*-min-mer corresponds to several locations in the genome, the resulting assembled sequence will be imperfect (similar to the output of `miniasm` [30]) which can be fixed by additional base-level polishing (not performed here).

#### How sequencing errors in base-space propagate to minimizer-space

In order to clarify the difference between base-space and minimizer-space in the presence of sequencing errors, we newly derive an expression of the expected error rate in minimizer-space (parameterized by *k,ℓ*, and *δ*), using a Poisson process model of random site mutations that was invoked by Mash [42]. Given the probability *d* of a single base substitution, the probability that no mutation will occur in a given *ℓ*-mer is *e^-ℓd^* under a Poisson model.

To estimate the number of erroneous *k*-min-mers in a read, we define for a given read *R*, the expected number *n_R_* of universe (*ℓ, δ*)-minimizers (described in the Preliminaries) in the read as *n_R_* = (|*R*| — *ℓ* +1) · *δ*. Since a *k*-min-mer is erroneous whenever at least one of *k* universe (*ℓ, δ*)-minimizers within the *k*-min-mer is erroneous, the probability that a given *k*-min-mer is erroneous is then 1 — *e^—ℓdk^*. The number of *k*-min-mers obtained from the read is *n_R_* — *k* + 1. Thus, the expected number of erroneous *k*-min-mers in a read is

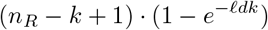

For instance, for a base-space mutation rate of *d* = 0.01, minimizer-space parameters *ℓ* = 12, *k* = 10, and *δ* = 0.01, and a read length of |*R*| = 20000, 70% of the *k*-min-mers in the read are erroneous. However, lowering the base-space mutation rate to *d* = 0.001 and keeping other values of *k* and *ℓ* identical renders only 10% of the *k*-min-mers erroneous within a read.

To estimate the average *ℓ*-mer identity of a read, we provide an approximation of the minimizer-space error rate given the base-space error rate. As seen above, an *ℓ*-mer that was selected as a universe minimizer has probability *e^-ℓd^* to be mutated. Mutations that occur outside of universe minimizers may now still affect the minimizer-space representation by turning a non-minimizer *ℓ*-mer into a universe minimizer (see Figure 2). Under the simplifying assumption that this effect occurs independently at each position in a read, the probability that an *ℓ*-mer turns into a universe minimizer is the probability of a mutation within that *ℓ*-mer times the probability *δ* that a random *ℓ*-mer is a universe minimizer, i.e. (1 — *e^-ℓd^*)*δ*. For a universe minimizer *m*, there are approximately 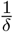 neighboring *ℓ*-mers that are candidates for turning into universe minimizers themselves due to a base error. We will conceptually attach those *ℓ*-mers to *m*, and consider that an error in any of those *ℓ*-mers leads to an insertion error next to *m*.

Combining the above terms leads to the following minimizer-space error rate approximation:

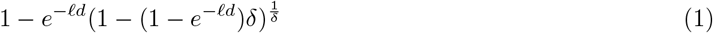

For an error rate of *d* = 5%, i.e. close to that of the Oxford Nanopore R10.3 chemistry, *ℓ* = 12, and *δ* = 0.01, the minimizer-space error rate is 65.1%, dropping to 2.3% when *d* = 0.1%. This analysis indicates that parameters *ℓ, k, δ* and the base error rate *d* together play an essential role in the performance of a mdBG-guided assembly.

#### Error correction using minimizer-space partial order alignment (POA)

Long-read sequencing technologies from Pacific Biosciences (PacBio) and Oxford Nanopore (ONT) recently enabled the production of genome assemblies with high contiguity, albeit with a relatively high error rate (≥ %5) in the reads, requiring either read error correction, and/or assembly polishing, both being resource-intensive steps [14, 36]. We will demonstrate that our minimizer-space representation is applicable to error-free sequencing reads and PacBio HiFi reads, which boast error rates lower than %1; however, in order to work with long reads with a higher error rate such as PacBio CLR and ONT, we present a resource-frugal error-correction step that uses partial order alignment (POA) [28], a graph representation of a multiple sequence alignment (MSA), in order to rapidly correct sequencing errors that occur in the minimizer-space representation of reads. Stand-alone error-correction modules such as `racon` [51] and `Nanopolish` [36] have also relied on POA for error-correction of long reads; however, these methods work in base-space, and as such, are still resource-intensive. We present an error-correction module that uses POA in minimizer-space that can correct errors in minimizer-space, requiring only the minimizer-space representation of reads as input.

An overview of the minimizer-space POA procedure is shown in Figure 3, and the detailed processes for the stages of the error-correction procedure are shown in Supplemental Note A. The input for the procedure is the collection of ordered lists of minimizers obtained from all reads in the dataset (one ordered list per read). As seen earlier, the ordered list of minimizers obtained from a read containing sequencing errors will likely differ from the one of an error-free read. However, provided the dataset has enough coverage, the content of other ordered lists of minimizers in the same genomic region can be used to correct errors in the query read in minimizer-space. To this end, we first perform a bucketing procedure for all ordered lists of minimizers using each of their *n*-tuples, where *n* is a user-specified parameter.

After bucketing, in order to initiate the error-correction of a query we collect its *neighbors*: Other ordered lists likely corresponding to the same genomic region. We use a distance metric (Jaccard or Mash [42] distance) to pick sufficiently similar neighbors. Once we obtain the final set of neighbors that will be used to error-correct the query, we run the partial order alignment (POA) procedure as described in [28], with the modification that a node in the POA graph is now a minimizer instead of an individual base, directed edges now represent whether two minimizers are adjacent in any of the neighbors, and edge weights represent the multiplicity of the edge in all of the neighbor ordered lists. After constructing the minimizer-space POA by aligning all neighbors to the graph, we generate a consensus (the best-supported traversal through the graph). Once the consensus is obtained in minimizer-space, we replace the query ordered list of minimizers with the consensus, and repeat until all reads are error-corrected. In order to recover the base-space sequence of the obtained consensus after POA, we store the sequence spanned by each pair of nodes in the edges, and generate the base-space consensus by concatenating the sequences stored in the edges of the consensus.

#### Implementation details

Reverse complementation is handled in our method in a natural way that is similar to classical base-space de Bruijn graphs. Each *ℓ*-mer is identified with its reverse complement, and a representative *canonical ℓ*-mer is chosen as the lexicographically smaller of the two alternatives. In turn, *k*-min-mers are identified with their reverse; no complementation is performed in minimizer-space as the complement of a canonical *ℓ*-mer is itself. Similarly to base-space assembly, any *k*-min-mer appearing only once in the multiset *V* is removed from *V* due to the likelihood that it is artefactual. Assembly graph simplifications are performed using `gfatools`^4^, with alternating rounds of tip clipping and bubble removal (Supplementary Note C), except for simulated perfect reads, which were only compacted into base-space unitigs.

In order to reduce memory usage, we write *k*-min-mers and the base-space sequences spanned by *k*-min-mers on disk, and retrieve them once the contigs are generated in minimizer-space. `rust-mdbg` includes a binary program (*to_basespace*) that transforms a simplified minimizer-space assembly into a base-space assembly.

#### Minimizer-space POA evaluation set-up

We extracted chromosome 4 (~ 1.2 Mbp) of the *D. melanogaster* reference genome, and simulated reads using the command `randomreads.sh pacbio=t of BBMap` [9]. We generated one dataset per error rate value from 0% to 10%, keeping other parameters identical (24 Kbp mean read length and 70X coverage). Reads were then assembled using our implementation with and without POA, using parameters *ℓ* = 10, *k* = 7, and *δ* = 0.0008 experimentally determined to yield a perfect assembly with error-free reads. We evaluated the average read identity in minimizer-space using semi-global Smith-Waterman alignment between the sequence of minimizers of a read and the sequence of minimizers of the reference, taking BLAST-like identity (number of minimizer matches divided by the number of alignment columns). We also evaluated the length of the longest reconstructed contig in base-space as a proxy for assembly quality.

## Results

An overview of our pipeline, implemented in Rust (`rust-mdbg`), is shown in Figure 4. We compared `rust-mdbg` to three recent assemblers optimized for low-error rate long reads: `Peregrine, HiCanu` [41] and `hifiasm` [10]. (See Supplementary Note D for versions and parameters.)

**Fig. 4:**
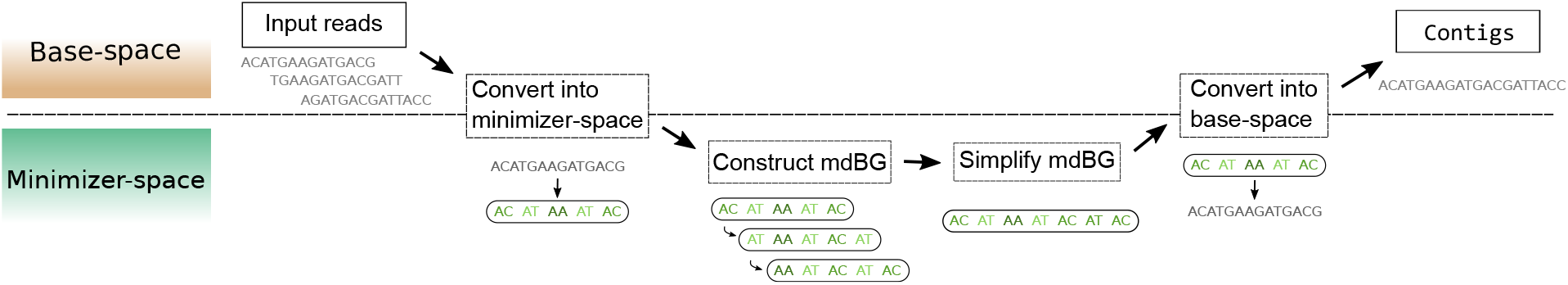
Overview of the assembly pipeline using mdBG. The region of the figure above (respectively below) the dotted line corresponds to analyses taking place in base-space (respectively minimizer-space). The input reads are converted into minimizer-space to construct a mdBG, which is then simplified and converted back into base-space to output contigs.

### Assembly using minimizer-space de Bruijn Graphs (mdBG)

#### Ultra-fast, memory-efficient and highly-contiguous assembly of real HiFi reads using rust-mdbg

We evaluated our software, `rust-mdbg`, on real Pacific Biosciences (PacBio) HiFi reads from *D. melanogaster*, at 100X coverage, and HiFi reads for human (HG002) at ~ 50X coverage, both taken from the HiCanu publication^5^ [41].

Since our method does not resolve both haplotypes in diploid organisms, we compared against the primary contigs of `HiCanu` and `hifiasm`. In our tests with *D. melanogaster*, the reference genome consists of all nuclear chromosomes from the RefSeq accession (GCA_000001215.4). Assembly evaluations were performed using `QUAST` [21] v5.0.2, run with parameters recommended in `HiCanu`’s article [41]. `QUAST` aligns contigs to a reference genome, allowing to compute contiguity and completeness statistics that are corrected for misassemblies (NGA50 and Genome fraction metrics respectively in Table 1). Assemblies were all run using 8 threads on a Xeon 2.60 GHz CPU. For `rust-mdbg` assemblies, contigs shorter than 50 Kbp were filtered out similarly to [41]. We did not report the running time of the base-space conversion step and graph simplifications as they are under 15% of the running CPU-time and run on a single thread, taking no more memory than the final assembly size, which is also less memory than the mdBG.

**Table 1:** Assembly statistics of *D. melanogaster* **real** HiFi reads (left), simulated perfect reads (center), and Human **real** HiFi reads (right), all evaluated using the commonly-used `QUAST` program. All assemblies were homopolymer-compressed. Wall-clock time is reported for 8 threads. NGA50 is a contiguity metric reported in Megabases (Mbp) by `QUAST` as the longest contig alignment to the reference genome so that shorter contig alignments collectively make up 50% of the genome length. The number of misassemblies is reported by `QUAST`. NGA50 and Genome fraction (Complete%) should be maximized, whereas all other metrics should be minimized. Only `Peregrine, hifiasm` and our method `rust-mdbg` were evaluated on Human assemblies, since HiCanu requires around an order of magnitude more running time.

| Tool | *D. mel* 100x real HiFi reads |  |  |  | *D. mel* 50x simulated perfect reads |  |  |  | Human real HiFi reads |  |  |
| --- | --- | --- | --- | --- | --- | --- | --- | --- | --- | --- | --- |
|  | Peregr. | HiCanu | hifiasm | rust-mdbg | Peregr. | HiCanu | hifiasm | rust-mdbg | Peregr. | hifiasm | rust-mdbg |
| Time | 40m11s | 7h43m | 5h17m | **1m9s** | 23m31s | 8h12m | 19h38m | **21s** | 14h8m | 58h41m | **10m23s** |
| Memory | 12 GB | 12 GB | 21 GB | **1.5 GB** | 16 GB | 18 GB | 51 GB | **<1 GB** | 188 GB | 195 GB | **10 GB** |
| # contigs | 682 | 928 | 538 | 93 | 63 | 45 | 48 | 34 | 8109 | 431 | 805 |
| NGA50(M) | 5.2 | 10.1 | 4.8 | 6.0 | 6.3 | 19.4 | 21.5 | 15.4 | 18.2 | 22.0 | 14.0 |
| Complete% | 93.9% | 96.6% | 96.6% | 90.8% | 98.2% | 98.1% | 98.2% | 96.2% | 97.0% | 94.2% | 95.5% |
| # misasm. | 10 | 5 | 0 | 0 | 3 | 5 | 0 | 1 | 312 | 7942 | 1073 |

Table 1 (leftmost) shows assembly statistics for *D. melanogaster* HiFi reads. Strikingly, `rust-mdbg` uses ~ 33x less wall-clock time and 8x less RAM than all other assemblers. In terms of assembly quality, all tools yielded high-quality results. `HiCanu` had 66% higher NGA50 statistics than `rust-mdbg`, at the cost of making more misassemblies, 385x longer runtime and 8x higher memory usage. `rust-mdbg` reported the lowest Genome fraction statistics, likely due in part to an aggressive tip-clipping graph simplification strategy, also removing true genomic sequences. To show the performance of lower-coverage assembly, additional results are presented in Supplementary Note E.

Table 1 (rightmost portion) shows assembly statistics for Human HiFi (HG002) reads. `rust-mdbg` performed assembly 81x faster with 18x less memory usage than `Peregrine`, at the cost of a 22% lower contiguity and 1.5% lower completeness, while making more misassemblies. Compared to `hifiasm, rust-mdbg` performed 338x faster with 19x lower memory, 36% lower contiguity and 1.3% higher completeness, making less misassemblies.

Remarkably, the initial unsimplified mdBG for the Human assembly only has around 12 million *k*-min-mers (seen at least twice in the reads, out of 40 million seen in total) and 24 million edges, which should be compared to the 2.2 Gbp length of the (homopolymer-compressed) assembly and the 100 GB total length of input reads in uncompressed FASTA format. This highlights that the mdBG allows very efficient storage and simplification operations over the initial assembly graph in minimizer-space.

#### Minimizer-space POA enables correction of reads with higher sequencing error rates

We introduce minimizer-space partial order alignment (POA) to tackle sequencing errors. To determine the efficacy of minimizer-space POA and the limits of minimizer-space de Bruijn graph assembly with higher read error rates, we performed experiments on a smaller dataset. In a nutshell, we simulated reads for a single *Drosophila* chromosome at various error rates and performed mdBG assembly with and without POA (see Methods for more details).

Figure 5 (left) shows that the original implementation without POA is only able to reconstruct the complete chromosome into a single contig up to error rates of 1%, after which the chromosome is assembled into ≥ 2 contigs. With POA, an accurate reconstruction as a single contig is obtained with error rates up to 4%. We further verified that, up to 3% error rate, the reconstructed contig corresponds structurally exactly to the reference, apart from the base errors in the reads. At 4% error rate, a single uncorrected indel in minimizer-space introduces a ~ 1 Kbp artificial insertion in the assembly.

**Fig. 5:**
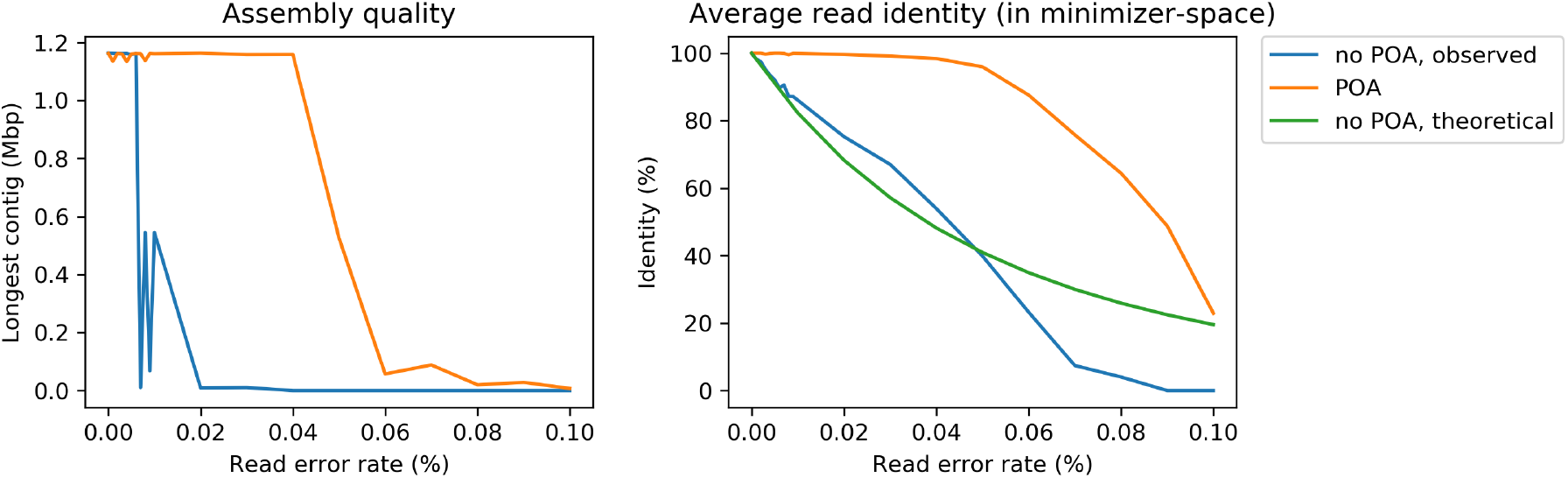
Effect of our minimizer-space POA correction on mdBG assembly. Reads from *D. melanogaster* chromosome 4 were simulated with base error rates ranging from 0%, 1%, …, up to 10%. Assemblies were run with and without minimizer-space POA correction. Left panel depicts the length of the longest contig for each assembly (uncorrected in blue, minimizer-space POA-corrected in orange). Right panel depicts the average read identity to the reference, computed in minimizer-space, for raw reads (observed in blue, and predicted by Equation 1 in green), and reads corrected by POA in minimizer-space (in orange).

Figure 5 (right) indicates that the minimizer-space identity of raw reads linearly decreases with increasing error rate. With POA, near-perfect correction can be achieved up to ~ 4% error rate, with a sharp decrease at > 5% error rates but still with an improvement in identity over uncorrected reads.

This highlights the importance of accurate POA correction: To put these results in perspective, mdBGs appear to be suitable to HiFi-grade data (< 1% error rates) without POA, and our POA implementation is almost, but not quite yet, able to cope with the error rate of ONT data (5%).

With POA, the runtime of our implementation was around 45 seconds and 0.4 GB of memory, compared to under 1 second and < 30 MB of memory without POA. Note that we did not use an optimized POA implementation; thus, we anticipate that further engineering efforts would significantly lower the runtime and possibly also improve the quality of correction.

#### Pangenome mdBG of a collection of 661K bacterial genomes allows efficient large-scale search of AMR genes

We applied mdBG to represent a recent collection of 661,405 assembled bacterial genomes [6]. To the best of our knowledge, this is the first de Bruijn graph construction of such a large collection of bacterial genomes. Previously only approximate sketches were created for this collection: A COBS index [5], allowing probabilistic membership queries of short *k*-mers (*k* = 31) [6], and sequence signatures (MinHash) using sourmash [45] and pp-sketch [29], none of which are graph representations.

The mdBG construction with parameters *k* = 10, *l* = 12, and *δ* = 0.001 took 3h50m wall-clock running time using 8 threads, totaling 8 hours CPU time (largely IO-bound). The memory consumption was 58 GB and the total disk usage was under 150 GB. Increasing *δ* to 0.01 yields a finer-resolution mdBG but increases the wall-clock running time to 13h30m, the memory usage to 481 GB, and the disk usage to 200 GB.

To compare the performance of mdBG with existing state-of-the-art tools for building de Bruijn graphs, we executed KMC 3 [25] to count 63-mers and Cuttlefish [24] to construct a de Bruijn graph from the counted *k*-mers. KMC 3 took 22 wall-clock hours and 191 GB memory using 8 threads, 2 TB of temporary disk usage, and 758 GB of output (56 billion distinct *k*-mers). Cuttlefish [24] did not terminate within three weeks of execution time. Hence, constructing the mdBG is at least two orders of magnitude more efficient in running time, and one order of magnitude in disk usage and memory usage.

Figure 6 shows the largest 5 connected components of the *δ* = 0.001 bacterial pangenome mdBG. As expected, several similar species are represented within each connected component. The entire graph consists of 16 million nodes and 45 million edges (5.3 GB compressed GFA), i.e. too large to be rendered, yet much smaller than the original sequences (1.4 TB lz4-compressed).

**Fig. 6:**
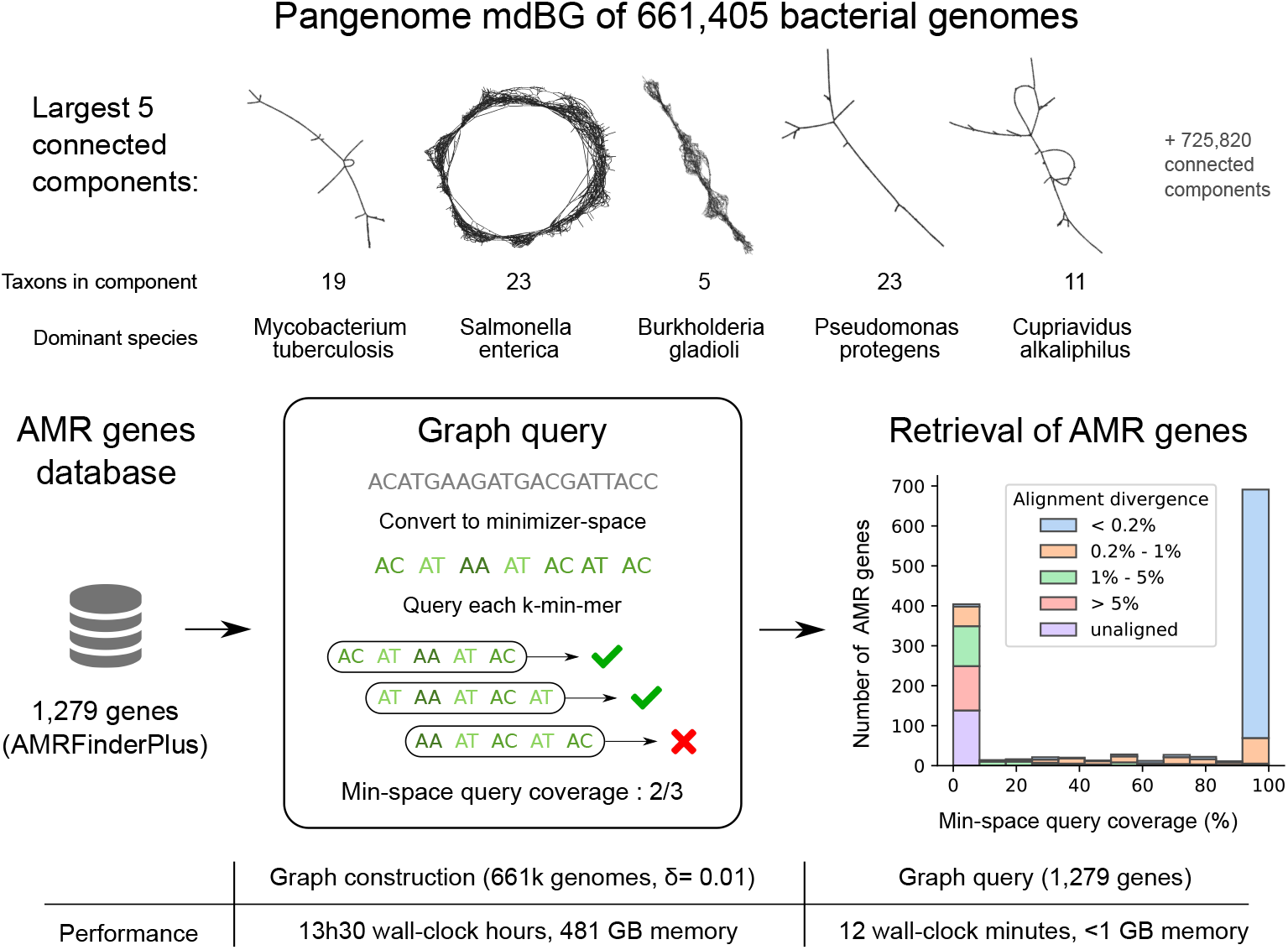
Pangenome mdBG of 661k bacterial genomes and retrieval of anti-microbial resistance genes. Top panel: first five connected components of the pangenome graph are displayed (using Gephi software). Each node is a *k*-min-mer, and edges are exact overlaps of *k* — 1 minimizers between *k*-min-mers. Middle panel: A collection of anti-microbial resistance gene targets was converted into minimizer-space, then each *k*-min-mer is queried in the 661k bacterial pangenome graph yielding a bimodal distribution of minimizer-space gene retrieval. The histogram is annotated by the minimal sequence diverge of each gene as aligned by `minimap2` to the pangenome over 90% of its length. Bottom panel: runtime and memory usage for each step. Note that the graph need only be constructed once in a preprocessing step.

To illustrate a possible application of this pangenome graph, we performed queries for the presence of AMR genes in the *δ* = 0.01 mdBG. We retrieved 1,502 targets from the NCBI AMRFinderPlus ‘core’ database (the whole `amr_targets.fa` file as of May 2021) and converted each gene into minimizer-space, using parameters *k* = 10, l = 12, *δ* = 0.01. Of these, 1,279 genes were long enough to have at least one *k*-min-mer (on average 10 *k*-min-mers per gene). Querying those *k*-min-mers on the mdBG, we successfully retrieved on average 61.2% of the *k*-min-mers per gene, however the retrieval distribution is bimodal: 53% of the genes have *ℓ* 99% *k*-min-mers found, and 31% of the genes have ≤ 10% *k*-min-mers found.

Further investigation of the genes missing from the mdBG was done by aligning the 661k genomes collection to the genes (in base-space) using `minimap2` (7 hours running time over 8 cores). We found that a significant portion of genes (141, 11%) could not be aligned to the collection. Also, *k*-min-mers of genes with aligned sequence divergence of 1% or more (267, 20%) did not match *k*-min-mers from the collection, and therefore had zero minimizer-space query coverage. Finally, although we performed sequence queries on a text representation of the pangenome graph, in principle the graph could be indexed in memory to enable instantaneous queries at the expense of higher memory usage.

This experiment illustrates the ability of mdBG to construct pangenomes larger than supported by any other method, and those pangenomes record biologically useful information such as AMR genes. Long sequences such as genes (containing at least 1 *k*-min-mer) can be quickly searched using *k*-min-mers as a proxy. There is nevertheless a trade-off of minimizer-space analysis that is akin to classical *k*-mer analysis: Graph construction and queries are extremely efficient however they do not capture sequence similarity below a certain identity threshold (in this experiment, around 99%). Yet, the ability of the mdBG to quickly enumerate which bacterial genomes possess any AMR gene with high similarity could provide a significant boost to AMR studies.

#### Highly-efficient assembly of real HiFi metagenomes using mdBG

We performed assembly of two real HiFi metagenome datasets (mock communities Zymo D6331 and ATCC MSA-1003, accessions SRX9569057 and SRX8173258). `Rust-mdbg` was run with the same parameters in the human genome assembly for the ATCC dataset, and with slightly tuned parameters for the Zymo dataset (Supplementary Note D).

Table 2 shows the results of `rust-mdbg` assemblies in comparison to `hifiasm-meta`, a metagenome-specific flavor of `hifiasm`. In a nutshell, `rust-mdbg` achieves roughly two orders of magnitude faster and more memory-efficient assemblies, while retaining similar completeness of the assembled genomes. Although Rust-mdbg metagenome assemblies are consistently more fragmented than `hifiasm-meta` assemblies, the ability of `rust-mdbg` to very quickly assemble a metagenome enables instant quality control and preliminary exploration of gene content of microbiomes at a fraction of the computing costs of current tools.

**Table 2:** Metagenome assembly statistics of the Zymo D6331 dataset (left) and the ATCC MSA-1003 dataset (right) using `hifiasm-meta` and `rust-mdbg`. The **Abundance** column shows the relative abundance of the species in the sample. The two rightmost columns show the species completeness of the assemblies as reported by `metaQUAST`. Table cells below 10% completeness are colored in red, below 98% in orange, and above in green.

| Zymo D6331 |  |  |  | ATCC MSA-1003 |  |  |  |
| --- | --- | --- | --- | --- | --- | --- | --- |
| Species | Abundance | hifiasm | rust-mdbg | Species | Abundance | hifiasm | rust-mdbg |
| *A. muciniphila* | 1.36% | 100.000% | 100.000% | *A. baumannii* | 0.18% | 99.839% | 99.955% |
| *B. fragilis* | 13.13% | 99.994% | 99.997% | *B. pacificus* | 1.80% | 100.000% | 99.998% |
| *B. adolescentis* | 1.34% | 100.000% | 99.730% | *B. vulgatus* | 0.02% | 81.846% | 70.895% |
| *C. albican* | 1.61% | 67.832% | 39.821% | *B. adolescentis* | 0.02% | 5.238% | 0.644% |
| *C. difficile* | 1.83% | 99.996% | 99.978% | *C. beijerinckii* | 1.80% | 99.993% | 99.993% |
| *C. perfringens* | 0.00% | 0.005% | 0.005% | *C. acnes* | 0.18% | 100.000% | 100.000% |
| *E. faecalis* | 0.00% | 0.006% | 0.006% | *D. radiodurans* | 0.02% | 82.499% | 53.659% |
| *E. coli B1109* | 8.44% | 100.000% | 97.918% | *E. faecalis* | 0.02% | 54.979% | 21.048% |
| *E. coli b2207* | 8.32% | 100.000% | 98.663% | *E. coli* | 18.0% | 100.000% | 100.000% |
| *E. coli B3008* | 8.25% | 100.000% | 99.558% | *H. pylori* | 0.18% | 100.000% | 100.000% |
| *E. coli B766* | 7.83% | 96.913% | 96.270% | *L. gasseri* | 0.18% | 97.779% | 98.142% |
| *E. coli JM109* | 8.37% | 100.000% | 97.852% | *N. meningitidis* | 0.18% | 98.593% | 99.030% |
| *F. prausnitzii* | 14.39% | 100.000% | 100.000% | *P. gingivalis* | 18.0% | 91.740% | 99.938% |
| *F. nucleatum* | 3.78% | 100.000% | 99.960% | *P. aeruginosa* | 1.80% | 99.706% | 99.726% |
| *L. fermentum* | 0.86% | 100.000% | 100.000% | *R. sphaeroides* | 18.0% | 99.748% | 100.000% |
| *M. smithii* | 0.04% | 99.840% | 87.175% | *S. odontolytica* | 0.02% | 8.179% | 1.046% |
| *P. corporis* | 5.37% | 99.561% | 99.561% | *S. aureus* | 1.80% | 100.000% | 100.000% |
| *R. hominis* | 3.88% | 100.000% | 100.000% | *S. epidermidis* | 18.0% | 100.000% | 100.000% |
| *S. cerevisiae* | 0.18% | 69.522% | 39.556% | *S. agalactiae* | 1.80% | 99.496% | 99.976% |
| *S. enterica* | 0.02% | 6.232% | 4.619% | *S. mutans* | 18.0% | 99.995% | 100.000% |
| *V. rogosae* | 11.02% | 100.00% | 100.000% |  |  |  |  |
| Running time |  | 34h29m | 55s | Running time |  | 59h16m | 3m51s |
| Memory usage |  | 83 GB | 0.9 GB | Memory usage |  | 313 GB | 1.3 GB |

## Discussion

Three areas we hope to tackle in our assembly implementation are: 1) its reliance on setting adequate assembly parameters; 2) lack of base-level polishing; and 3) haplotype separation. Regarding 1), we are experimenting with automatic selection of parameters *ℓ, k* and *δ*. A heuristic formula is presented along with its implementation and results in the GitHub repository of `rust-mdbg`; however, it leads to lower-quality results (e.g. 1 Mbp N50 for the HG002 assembly versus 14 Mbp in Table 1). We also provide a preliminary multi-k assembly script inspired by IDBA [43]. While automatically setting mdBG parameters is fundamentally a more complex task than just determining a single parameter (`k`) in classical de Bruijn graphs, we anticipate that similar techniques to `KmerGenie` [13] could be applicable, where optimal values of (*ℓ, k, δ*) would be found as a function of the *k*-min-mer abundance histogram.

Regarding directions 2) and 3), polishing could be performed as an additional step, by feeding the reads and the unpolished assembly to a base-space polishing tool such as `racon` [51]. Haplotype separation might prove more difficult to incorporate in mdBGs: Unlike HiFi assemblers which use overlap graphs with near-perfect overlaps, minimizer-space de Bruijn graphs cannot differentiate between exact and inexact overlaps in bases that are not captured by a minimizer. However, an immediate workaround is to perform haplotype phasing on resulting contigs, using tools such as `HapCut2` [19] or `HapTree-X` [4].

We anticipate that *k*-min-mers could become a drop-in replacement for ubiquitously-adopted *k*-mers for the comparison and indexing of long, highly similar sequences, e.g. in genome assembly, transcriptome assembly, and taxonomic profiling.

### Key Resources Table

| Resource | Source | Identifier |
| --- | --- | --- |
| **Deposited Data** |  |  |
| *D. melanogaster* and *H. sapiens* HiFi reads | [41] | Table 1 [https://doi.org/10.1101/gr.263566.120] |
| ATCC MSA-1003 and Zymo D6331 HiFi reads |  | Table 2 [SRA identifiers SRX9569057 and SRX8173258] |
| **Software and Algorithms** |  |  |
| mDBG code | This paper | https://github.com/ekimb/rust-mdbg/ |

## Acknowledgements

R.C. was funded by ANR Inception (ANR-16-CONV-0005) and PRAIRIE (ANR-19-P3IA-0001) grants. B.B. and B.E. were funded by the NIH R01HG010959 grant. The authors are grateful to A. Limasset and B. Hie for remarks on the manuscript.

## Supplementary Information

### A Minimizer-space POA additional methods

#### POA bucketing and preprocessing

In Algorithm 1, all tuples of length *n* of an ordered list of minimizers are computed using a sliding window (lines 4-6), and the ordered list of minimizers itself is stored in the buckets labeled by each *n*-tuple (line 7). We use bucketing as a proxy for set similarity, since each pair of reads in the same bucket will have an *n*-tuple (the label of the bucket), and will be more likely to come from the same genomic region.

##### Algorithm 1 Bucketing procedure for all ordered lists of minimizers

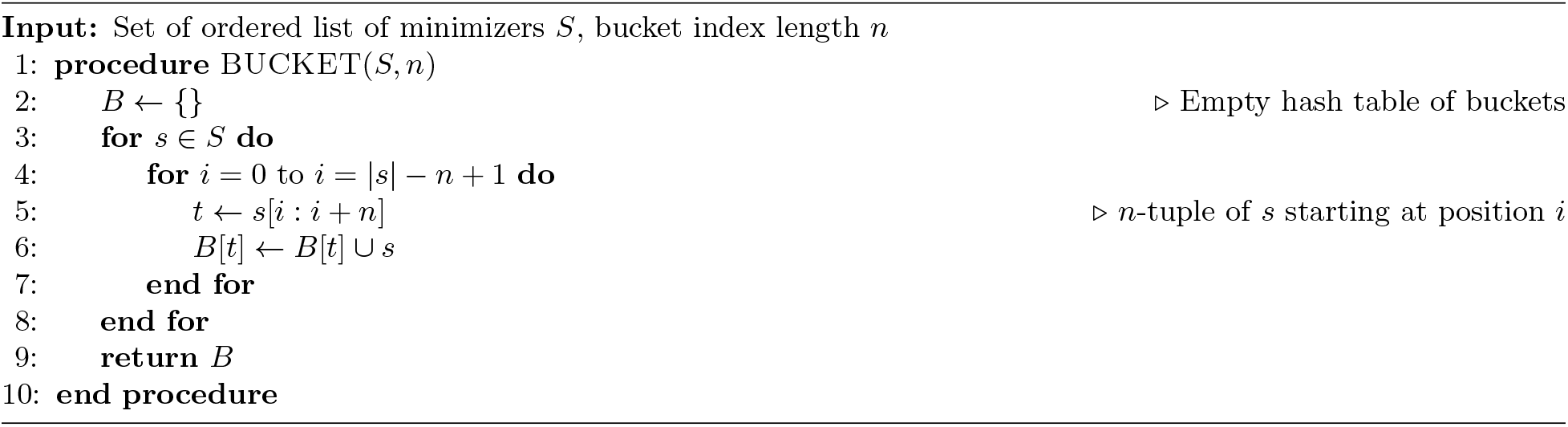

The overview of the collection of neighbors for error-correcting a query ordered list of minimizers is shown in Algorithm 2. We obtain all *n*-tuples of a query ordered list, and collect the ordered lists in the previously populated buckets indexed by its *n*-tuples (lines 10-15). These ordered lists are viable candidates for neighbors, since they share a tuple of length at least *n* with the query ordered list; however, since a query *n*-tuple may not uniquely identify a genomic region, we apply a similarity filter to further eliminate candidates unrelated to the query. Using either Jaccard or Mash distance [42] as a similarity metric, for a user-specified threshold *φ*, we filter out all candidates that have distance ≥ *φ* to the query ordered list to obtain the final set of neighbors that will be used for error-correcting the query (lines 1-9).

#### POA graph construction and consensus generation

Algorithm 4 describes a canonical POA consensus generation procedure, similar to `racon` [51], except that here consensus is performed in minimizer-space.

The minimizer-space POA error-correction procedure is shown in Algorithm 3. For each neighbor of the query, we perform semi-global alignment between a neighbor ordered list and the graph, where for two minimizers *m_i_* and *m_j_*, a match is defined as *m_i_* = *m_j_*, and a mismatch is defined as *m_i_* ≠ *m_j_* (lines 17-19). After building the POA graph *G* = (*V, E*) by aligning all neighbors in minimizer space, we generate a consensus to obtain the best-supported traversal through the graph. We first initialize a scoring λ, and set λ[*v*] = 0 for all *v* ∈ *V*. Then, we perform a topological sort of the nodes in the graph, and iterate through the sorted nodes. For each node v, we select the highest-weighted incoming edge *e* = (*u,v*) with weight *w_e_*, and set λ[*v*] = *w_e_* + λ(*u*). The node *u* is then marked as a predecessor of *v* (lines 21-28).

## B Exploration of rust-mdbg parameter space on simulated perfect reads

In order to demonstrate the efficacy of our approach in terms of results quality in an ideal setting, we simulated error-free reads of length 100 Kbp at 50X coverage of the *D. melanogaster* genome. The parameters for the assembly were *k* = 30, *ℓ* = 12, and *δ* = 0.0005. Table 1 (center) shows that `rust-mdbg` is able to assemble these error-free reads nearly as well as `HiCanu` and `hifiasm`, within lower but similar NGA50 (~ 25% lower) and genome fraction (< 1% lower) values. However, `rust-mdbg` is 2-3 orders of magnitudes faster and uses an order of magnitude less memory.

### Algorithm 2 Collection of neighbors for a given query ordered list

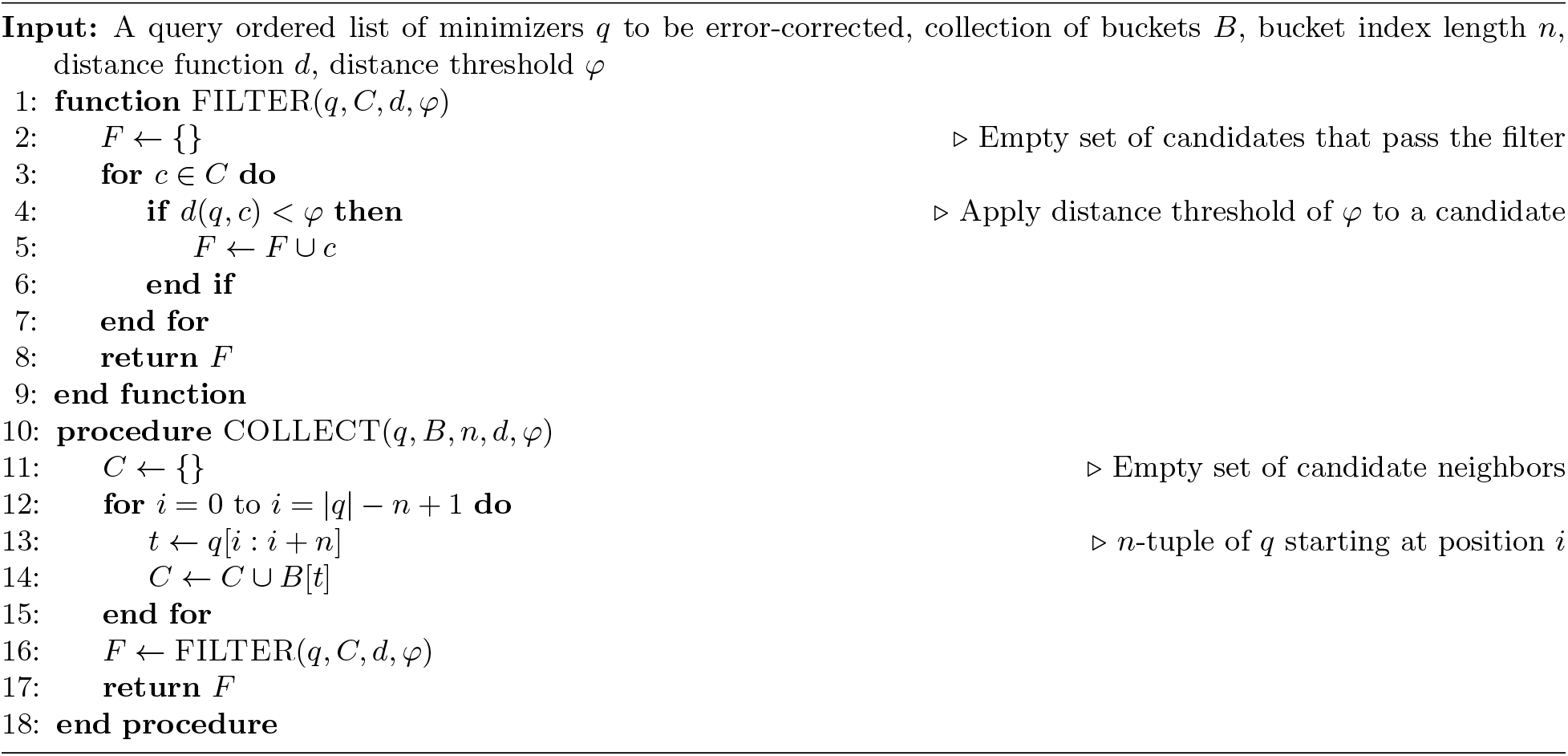

For a base-space de Bruijn graph assembler, the quality of the assembly depends on a single parameter (*k*), whereas in a `rust-mdbg` assembly, there are three parameters (*ℓ, k, δ*) that can affect assembly quality independently (see Methods). We investigated the effect of changing *k* for given *ℓ* and *δ*, and changing *δ* for given *k* and *ℓ* on the performance of `rust-mdbg` on perfect reads. For *ℓ* = 12 and *k* = 30, we tested different values for *δ* from 0.0001 to 0.0005 (increased by 0.00005 in each iteration). For *ℓ* = 12 and *δ* = 0.0003, we tested different values of *k* from 10 to 50 (increased by 1 in each iteration). For each iteration, we computed the *k*-min-mer recovery rate (the percentage of *k*-min-mers obtained from the reads that also exist in the set of *k*-min-mers from the reference) as a means of quantifying the quality of a minimizer-space assembly through a completeness metric.

Figure S1 shows the results of this investigation. For fixed values of *k* = 30 and *ℓ* = 12, *k*-min-mer recovery rate is insufficiently low for *δ* < 0.00025: Since the ordered lists of minimizers obtained from the reads need to have length > *k* in order to not be discarded, a very low density value causes a higher fraction of reads to be skipped, decreasing *k*-min-mer recovery rate. For *δ* ≥ 0.00025, an increasingly smaller portion of the reads are discarded, consistently yielding *k*-min-mer recovery rates of > 90%. We further observe that for fixed values of *δ* = 0.0003 and *ℓ* = 12, *k*-min-mer recovery rate is consistently above 95% for *k*-min-mer lengths of 10 to 35. Since *δ* = 0.0003, a sufficient portion of the reads are transformed into *k*-min-mers at this *k*-min-mer length, and higher values of *k* will result in a larger portion of the reads to be discarded.

## C gfatools command lines

The following (relatively aggressive) GFA assembly graph simplifications rounds were performed for all mdBG assemblies, using https://github.com/lh3/gfatools/. Rounds are of two types: `-t x,y` removes tips having at most *x* segments and of maximal length *y* bp, and `-b z` remove bubbles of maximal radius *z* bp. In addition, `gfa_break_loops.py` is a custom script (available in the `rust-mdbg` GitHub repository) that removes self-loops in the assembly graph, as well as an arbitrary edge in *x* ↔ *y* cycles.

~~~
gfatools asm -t 10,50000 -t 10,50000 -b 100000 -b 100000 -t 10,50000 \
             -b 100000 -b 100000 -b 100000 -t 10,50000 -b 100000 \
             -t 10,50000 -b 1000000 -t 10,150000 -b 1000000 -u > $base.tmp1.gfa
gfa_break_loops.py $base.tmp1.gfa > $base.tmp2.gfa
gfatools asm $base.tmp1.gfa -t 10,50000 -b 100000 -t 10,100000 \
             -b 1000000 -t 10,150000 -b 1000000 -u > $base.tmp3.gfa
gfa_break_loops.py $base.tmp3.gfa > $base.tmp4.gfa
gfatools asm $base.tmp4.gfa -t 10,50000 -b 100000 -t 10,100000 \
             -b 1000000 -t 10,200000 -b 1000000 -u > $base.msimpl.gfa
~~~

### Algorithm 3 Minimizer-space POA graph construction and consensus generation

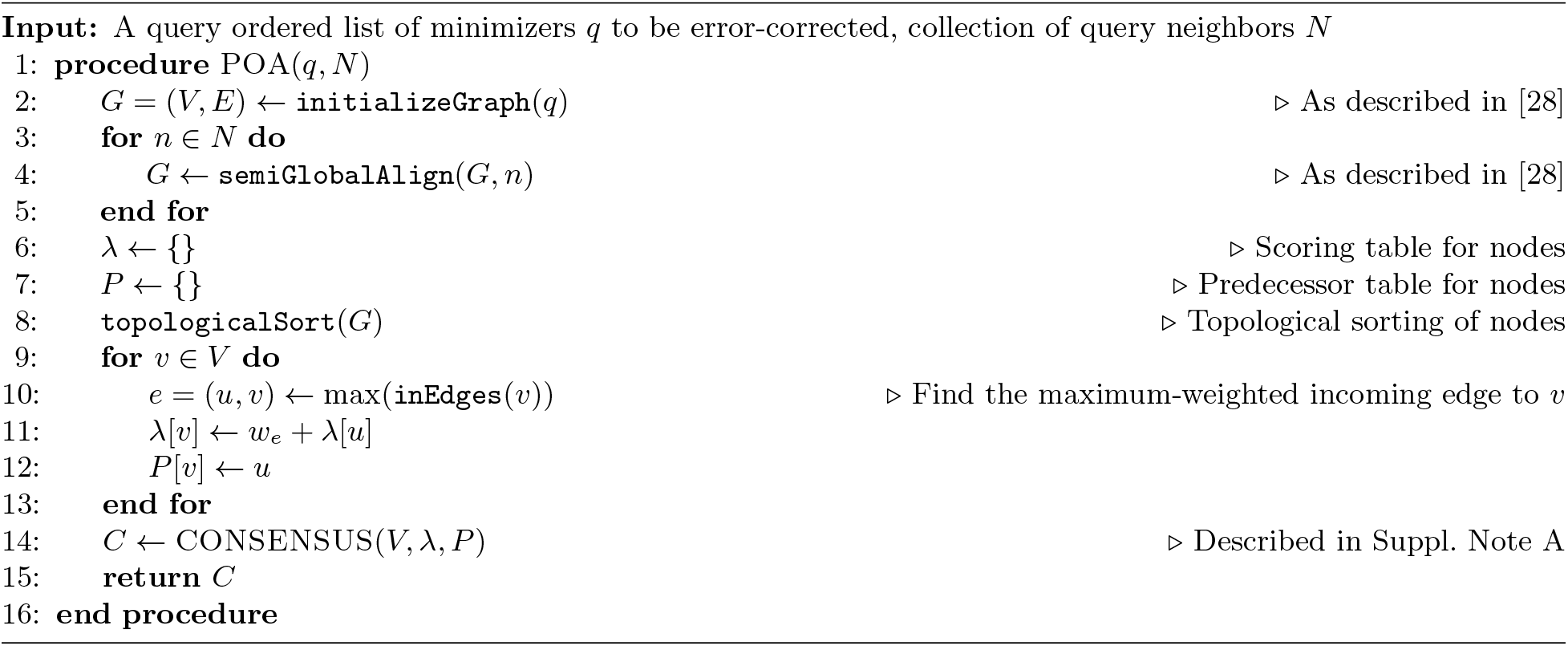

### Algorithm 4 Consensus generation on POA graph

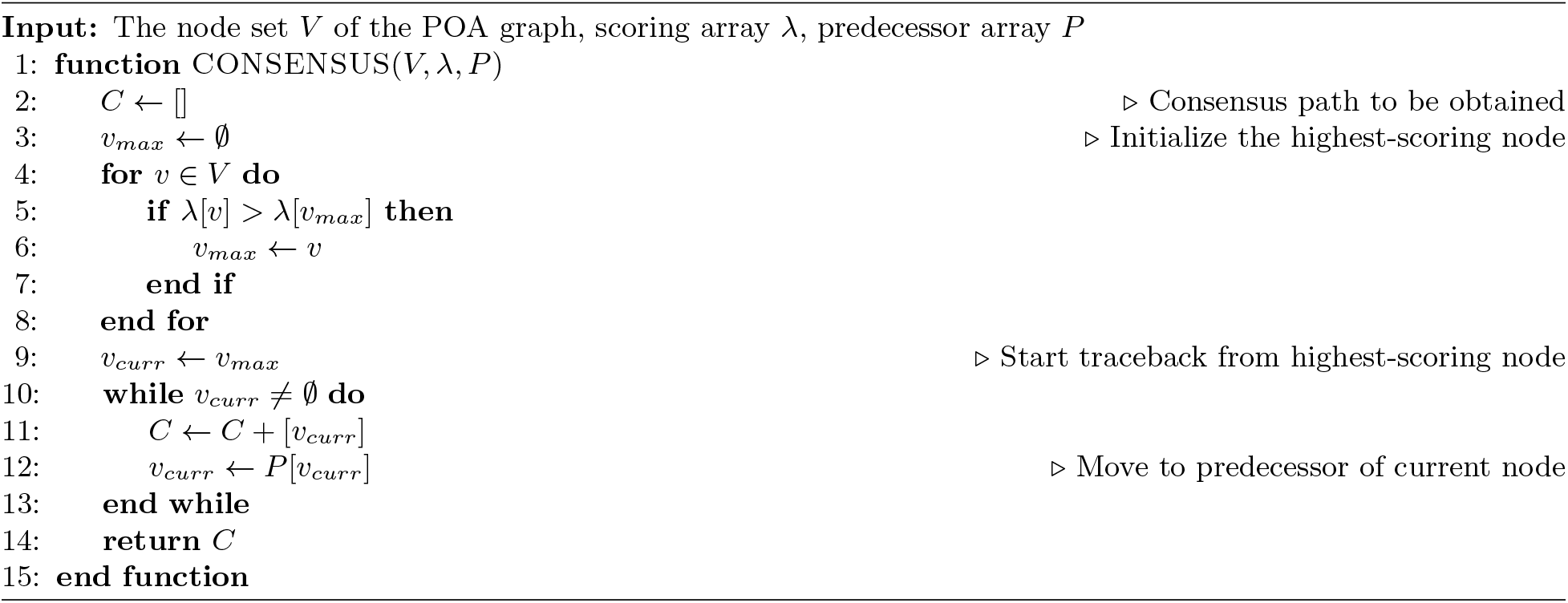

## D Genome assembly tools, versions and parameters

`HiCanu` (v2.1) was ran with default parameters, `hifiasm` (commit `8cb131d`) with parameters `-10 -f0`, and `Peregrine` (commit `008082a`) with command line: `8 8 8 8 8 8 8 8 8 --with-consensus --shimmer-r 3 --best_n_ovlp 8. rust-mdbg` was run with parameters *k* = 35, *ℓ* = 12, and *δ* = 0.002 for *D. melanogaster*, and *k* = 21, *ℓ* = 14, *δ* = 0.003 for HG002.

**Fig. S1:**
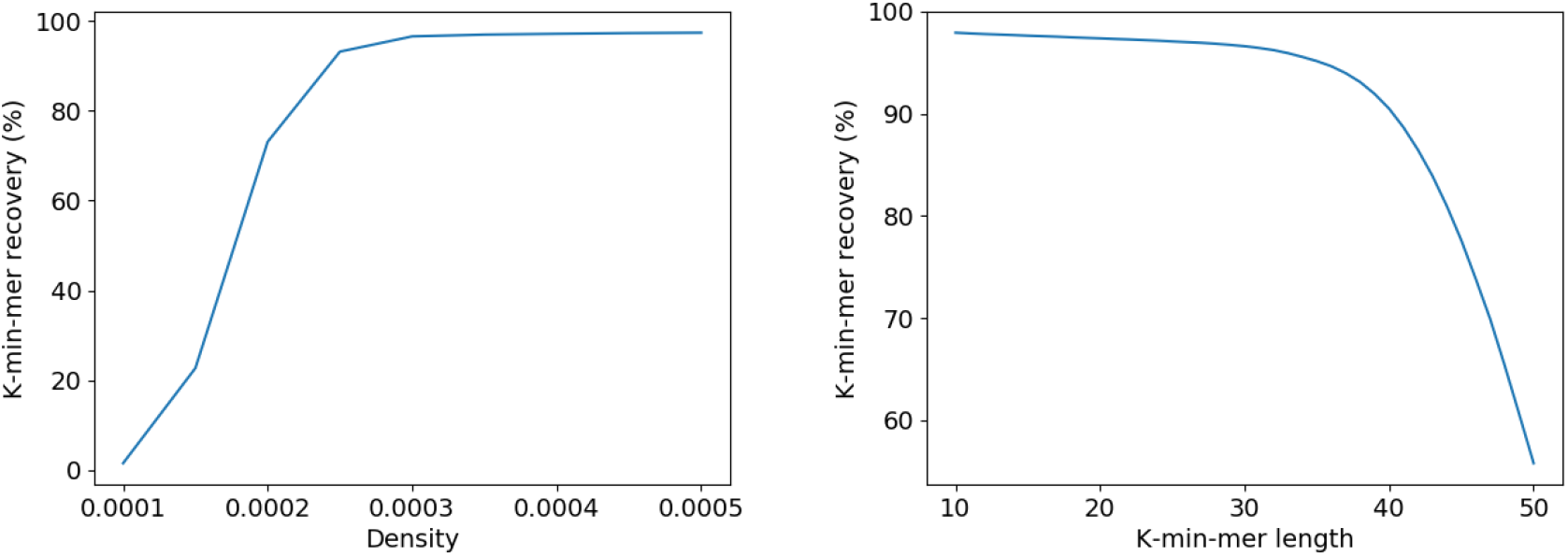
Robustness of rust-mdbg assemblies by varying the *k* and *δ* parameters, on *D. melanogaster* simulated perfect reads. The proportion of recovered *k*-min-mer values is reported in both plots. Left panel shows recovery rates for *k* = 30, *ℓ* = 12, and varying *δ* from 0.0001 to 0.0005. Right panel shows recovery rates for *ℓ* = 12, *δ* = 0.0003 and varying *k* from 10 to 50.

For metagenomes, `rust-mdbg` was run with parameters *k* = 21, *ℓ* = 14, *δ* = 0.003 for the ATCC MSA-1003 dataset (same parameters as the human dataset), and *k* = 40, *ℓ* = 12, *δ* = 0.004 for the Zymo D6331 dataset. `Hifiasm-meta` (commit `cda13b8`) was run with parameters `-S --lowq-10 50` for ATCC MSA-1003 and default for Zymo.

## E Lower-coverage assembly of *D. melanogaster* HiFi reads

Table S1 presents an assembly of 40X coverage of *D. melanogaster* reads sampled from the `HiCanu` publication^6^ [41].

**Table S1:** **Assembly statistics** of *D. melanogaster*, real HiFi 40X reads, evaluated using the same metrics as in Table 1. For `rust-mdbg`, parameters were *k* = 35, *ℓ* = 12, and *δ* = 0.002.

| Tool name | *D. mel* 40X real HiFi reads |  |  |  |
| --- | --- | --- | --- | --- |
|  | Peregrine | HiCanu | hifiasm | mdBG |
| Wall-clock time (8 threads) | **56m15s** | **5h29m** | **2h24m** | **35s** |
| Memory usage | 7 GB | 16 GB | 9 GB | 1 GB |
| Number of contigs | 516 | 99 | 157 | 125 |
| NGA50 (Mbp) | 8.3 | 10.0 | 15.3 | 2.1 |
| Genome fraction (%) | 94.3 | 90.4 | 94.4 | 84.0 |
| # of misassemblies | 4 | 5 | 3 | 0 |

## F Locally Consistent Parsing (LCP)

Locally Consistent Parsing (LCP) describes sets of evenly spaced *core substrings* of a given length *ℓ* that cover any string of length *n* for any alphabet [16]. The set of core substrings can be pre-computed such that a string of length *n* is covered by ~ *n/ℓ* core substrings on average. LCP and the concept of core substrings were used in the first linear-time algorithm for approximate string matching [16], for string indexing under block edit distance [39], and for almost linear-time approximate string alignment [1].

SCALCE [22] introduced LCP to genome compression, and used the longest core substring(s) in each read as representatives to group together similar reads, which are then reordered lexicographically for compression without the need of a reference genome. In preliminary testing of LCPs as an alternative to minimizers in our pipeline, we integrated the pre-computed set of core substrings described in SCALCE into the universe (*ℓ, δ*)-minimizers scheme in `rust-mdbg`, where we selected an *ℓ*-mer *m* as a minimizer if *m* is a universe (*ℓ, δ*)-minimizer and also appears in the set of core substrings. We evaluated both minimizer schemes on simulated perfect reads from *D. melanogaster* at 50X coverage, real Pacific Biosciences HiFi reads from *D. melanogaster* at 100X coverage, and HiFi reads for human (HG002) at ~ 50X coverage, taken from the `HiCanu` publication^7^ [41]. We did not notice a major difference using LCP versus only universe minimizers, but our implementation should be seen as a baseline for future optimizations.

**Table S2.** **Assembly statistics** using both universe minimizers (grey, same results as in Table 1) and universe minimizers with LCP (black) of *D. melanogaster* real HiFi reads (left), simulated perfect reads (center), and Human real HiFi reads (right), evaluated using the same metrics in Table 1. Parameters for both schemes were *k* = 35, *ℓ* = 12, and *δ* = 0.002 for *D. melanogaster*, and *k* = 21, *ℓ* = 14, and *δ* = 0.003 for Human.

| Minimizers scheme | *D. mel* 100X real HiFi reads |  | *D. mel* 50X simulated perfect reads |  | Human real HiFi reads |  |
| --- | --- | --- | --- | --- | --- | --- |
|  | Universe | Universe + LCP | Universe | Universe + LCP | Universe | Universe + LCP |
| Time | 46s | 48s | 21s | 22s | 10m23s | 10m31s |
| Memory | 1 GB | 1 GB | < 1 GB | < 1 GB | 10 GB | 10 GB |
| # contigs | 109 | 106 | 34 | 35 | 805 | 807 |
| NGA50(M) | 5.5 | 5.4 | 15.4 | 15.4 | 14.0 | 13.9 |
| Complete% | 83.2% | 83.3% | 96.2% | 96.3% | 95.5% | 95.5% |
| # misasm. | 0 | 0 | 1 | 2 | 1073 | 1139 |

## Footnotes

4 https://github.com/lh3/gfatools

5 https://obj.umiacs.umd.edu/marbl_publications/hicanu/index.html

6 https://obj.umiacs.umd.edu/marbl_publications/hicanu/index.html

7 https://obj.umiacs.umd.edu/marbl_publications/hicanu/index.html

